# DataSAIL: Data Splitting Against Information Leakage

**DOI:** 10.1101/2023.11.15.566305

**Authors:** Roman Joeres, David B. Blumenthal, Olga V. Kalinina

## Abstract

Information Leakage is an increasing problem in machine learning research. It is a common practice to report models with benchmarks, comparing them to the state-of-the-art performance on the test splits of datasets. If two or more dataset splits contain identical or highly similar samples, a model risks simply memorizing them, and hence, the true performance is overestimated, which is one form of Information Leakage. Depending on the application of the model, the challenge is to find splits that minimize the similarity between data points in any two splits. Frequently, after reducing the similarity between training and test sets, one sees a considerable drop in performance, which is a signal of removed Information Leakage. Recent work has shown that Information Leakage is an emerging problem in model performance assessment.

This work presents DataSAIL, a tool for splitting biological datasets while minimizing Information Leakage in different settings. This is done by splitting the dataset such that the total similarity of any two samples in different splits is minimized. To this end, we formulate data splitting as a Binary Linear Program (BLP) following the rules of Disciplined Quasi-Convex Programming (DQCP) and optimize a solution. DataSAIL can split one-dimensional data, e.g., for property prediction, and two-dimensional data, e.g., data organized as a matrix of binding affinities between two sets of molecules, accounting for similarities along each dimension and missing values. We compute splits of the MoleculeNet benchmarks using DeepChem, the LoHi splitter, GraphPart, and DataSAIL to compare their computational speed and quality. We show that DataSAIL can impose more complex learning tasks on machine learning models and allows for a better assessment of how well the model generalizes beyond the data presented during training.

## 1 Introduction

Machine learning (ML) is one of the fastest-growing fields in computer science, leading to advances in many computer and life sciences domains, including bioinformatics. The most prominent examples are DeepMind’s AlphaFold[1] for predicting protein three-dimensional structure, introducing similar models, such as OmegaFold[2] and OpenFold[3]. Another rapidly developing field of research is protein language models, which compute high-dimensional embeddings of proteins based on millions of sequences [4], which can be subsequently used in a variety of downstream tasks, ranging from predicting protein secondary structure to estimating their luminescence [5]. Other bioinformatics fields also benefit from using (simpler) machine learning models [6].

The sub-field of machine learning most relevant for this study is *Supervised Learning*, where the general approach to solving a problem can be described as follows: Given the dataset 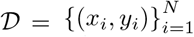 of *N* features *x*_*i*_ ∈*X* and labels *y*_*i*_ ∈*Y*, learn a function *f*_*θ*_ : *X* → *Y* that minimizes a loss function *ℒ* (*f*_*θ*_(*x*_*i*_), *y*_*i*_). This is achieved by adjusting the function’s learnable parameters *θ* and selecting a function *f* ∈*ℋ* from the hypothesis space [7, 8]. The development of a model has two levels: The hyperparameters, i.e., selecting *f* from *ℋ*, and the parameters *θ* of *f* . Therefore, one needs three pairwise disjoint datasets to (i) train the parameters *θ*, (ii) optimize hyperparameters *f* (these can include architecture, learning rate, or optimizer), and (iii) assess the performance of the model on so far unseen data. For this, the dataset 𝒟 is split into pairwise disjoint subsets 𝒟_*train*_, 𝒟 _*val*_, and 𝒟_*test*_ . It is important to use unseen data to check the ability of the model to generalize to unseen samples. Suppose the samples for performance estimation are too similar to the training samples and are not representative of the data the model will see during inference. In that case, the model might “cheat” and perform well just by memorizing properties instead of generalizing based on features. This phenomenon is called Information Leakage or Data Leakage [9]. Recent studies show that Information Leakage is an emerging problem in many fields, leading to inflated performances and overoptimistic conclusions [10]. Whalen et al. identify Information Leakage as one of the five most pressing pitfalls when applying ML in genomics [11]. For example, firm evidence exists that current models for predicting protein-protein interactions are plagued by Information Leakage. When it is removed, they perform no better than random guessing [12, 13, 14].

DataSAIL is focused on splitting biologically relevant datasets of molecules, for example, for property or interaction prediction tasks. Formally, we call a dataset in which one output value (measurement) corresponds to one molecule a *one-dimensional* dataset (see Figure 1A). An example of such a dataset is a set of small molecules with measured solubility. If an output measurement corresponds to two molecules, for instance, in the case of drug-target interactions, we call such data a *two-dimensional* dataset (Figure 1). Notably, the similarity between molecules can be defined over each dimension in a two-dimensional dataset.

**Figure 1:**
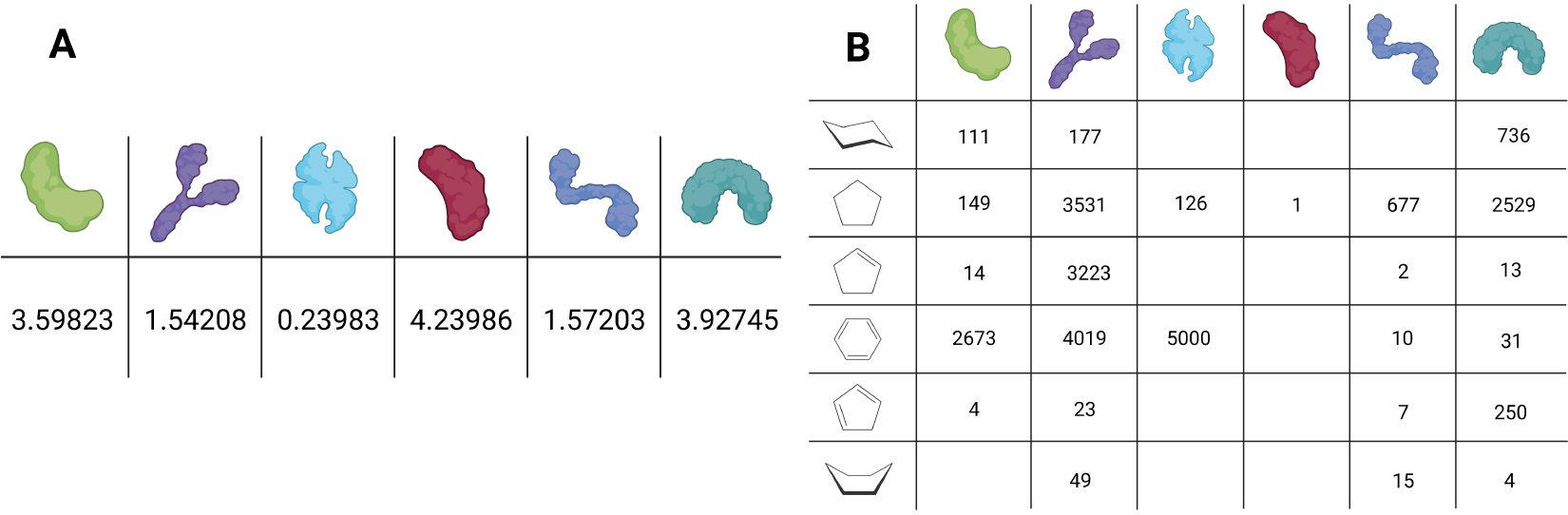
Visualization of one-dimensional data (in A) and two-dimensional data (in B).

One source of Information Leakage is having the same samples in multiple splits. We will call this source *I*, as it is based on the identities of the data points. This is usually easy to control for, and this is done routinely. Another source is having (too) similar samples in two splits. An example of this in a task of molecular property prediction can be very similar molecules, e.g., paracetamol and phenacetin, that naturally have very similar properties. If many such examples are present in the dataset, a model might achieve a good performance estimate by memorizing the data instead of generalizing meaningful features. This source of Information Leakage will be called *C* as it can be defined over clusters in the data. This is usually more difficult to control. One can split the data based on the sample identifiers to prevent Information Leakage caused by *I*. This is called cold-drug, cold-target, or cold-protein split in the drug-target interaction setting. But to reduce Information Leakage by *C*, one needs to cluster the data before assigning the clusters to splits. There are packages solving *I* and partially tackling *C*, e.g. TDCommons[15], DeepChem[16], LoHi-Splitter[17], and GraphPart[18]. Section 1.4 discusses those algorithms in more detail.

We define different splitting tasks with abbreviations based on the source of Information Leakage that is tackled (*I* or *C*) and the dimensions of the dataset. Those tasks include random split (*R*), identity-based one-dimensional cold split (*I1*), identity-based two-dimensional cold split (*I2*), cluster-based one-dimensional cold split (*C1*), and cluster-based two-dimensional cold split (*C2*). Figure 2 gives a visualization of those for one-dimensional and two-dimensional datasets. Assigning every interaction to a split in two-dimensional cold splits cannot be guaranteed. The reason is that the two interacting data points can be assigned to different splits. Therefore, their interaction is across splits; one cannot assign it to any split without leaking information.

**Figure 2:**
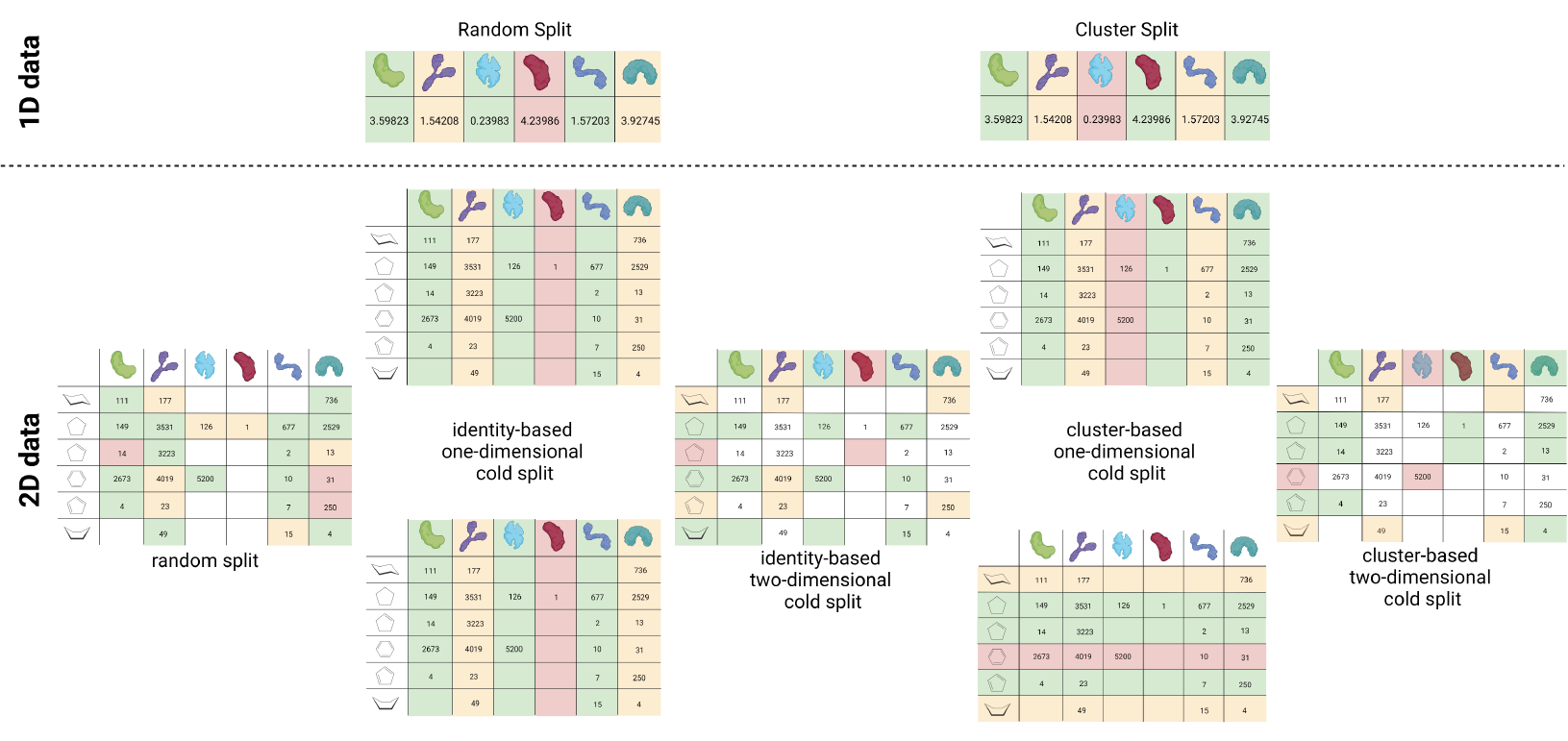
Visualization of the different splitting techniques based on exemplary datasets from Figure 1.Furthermore, the connection between one-dimensional and two-dimensional data splits is shown. Tiles with a green background are samples assigned to training, yellow are validation samples, and the test set is red. Tiles that are not assigned are left white.

In this work, we present DataSAIL, a Python package that implements splitting based on data point identities and clusters of similar data points, emphasizing applications for biological macromolecules and chemicals. DataSAIL improves the quality of data splits while reducing the runtime to compute them. This will be demonstrated by training D-MPNN[19] and DeepDTA[20] on splits of MoleculeNet benchmarks[21] computed with DataSAIL, DeepChem[22], GraphPart [18], and the LoHi-Splitter[17].

### 1.1 Clustering vs. Splitting

Before diving into the mathematical details, we discuss some terminology that will help us to precisely describe the idea of DataSAIL. According to Madhulatha, “*clustering* is the process of grouping similar objects into different groups, or more precisely, the partitioning of a data set into subsets so that the data in each subset according to some defined distance measure” [23, p.1]. The term *splitting* is often used interchangeably with clustering, which is unacceptable for this work. We refer to *splitting* when discussing partitioning a dataset into a fixed number of sets later used to train, validate, and test a model. When splitting a dataset, the computed partitions are called *splits*.

Splitting and clustering are very similar tasks as both try to find substructures in a dataset and partition them based on the substructures (Figure 3). The main difference between clustering and splitting lies in the goal of each procedure. Clustering is widely used in data analysis and aims to group samples in a dataset based on their similarity or distance from each other. Splitting, on the other hand, is primarily used in the machine-learning context. Splitting aims to divide the dataset into a fixed number of fixed-size partitions. Whereas in clustering, the defining parameter is the number of clusters or distance between them, in splitting, it is essential to maintain a specific ratio of data points between the splits for training purposes.

**Figure 3:**
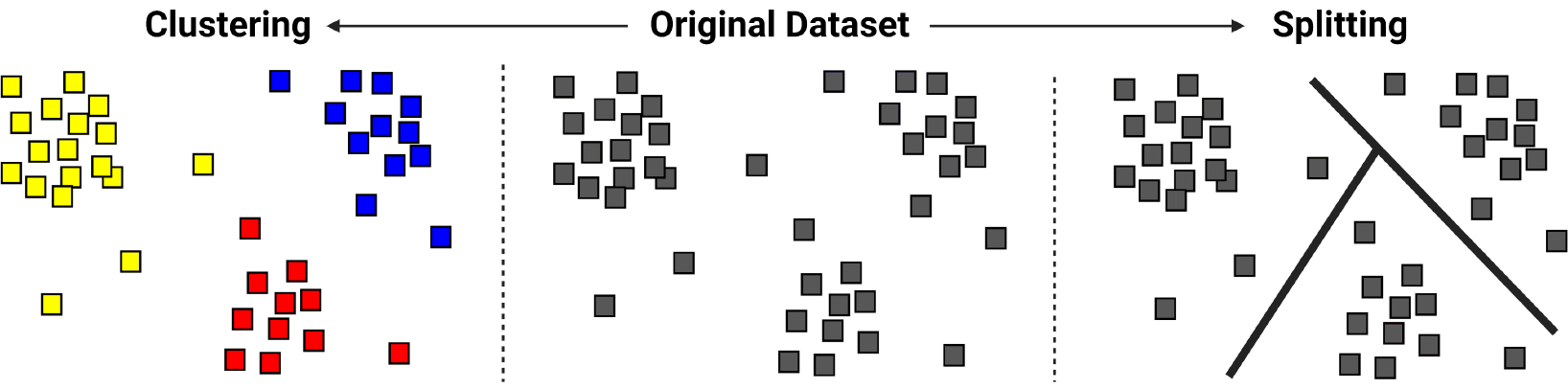
Visualization of the difference between clustering and splitting.

### 1.2 Disciplined Quasi-Convex Programming

In this section, we introduce Disciplined Quasi-Convex Programming(DQCP) by considering each term separately.

#### 1.2.1 Mathematical Programming

A mathematical program is an optimization problem that has the following form or can be converted into that form:

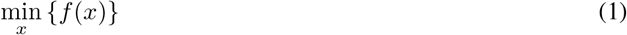

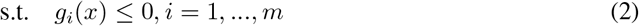

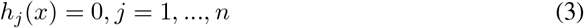

Equation 1 defines the objective function *f* to be minimized by finding the optimal value for the variable *x*. The adjustable variables of a mathematical program are indicated under the min operator. Equations 2 and 3 define *m* inequalities and *n* equalities that must be fulfilled while finding the optimal value. If *m* = *n* = 0, the problem is called unconstrained; otherwise, the constraints *g*_*i*_ and *h*_*i*_ define bounds for *x*, within which *x* is optimized [24].

#### 1.2.2 Convex and Quasi-Convex Functions

Let *C* be a convex subset of a real vector space. For every convex function *f* : *C*→ ℝ with *C* ⊆ ℝ^*n*^ for any *n*,

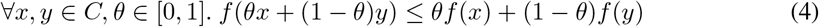

holds. The formula expresses that for any two points *x* and *y* in the domain of *f* the function values of *f* between *x* and *y*, i.e., *f* (*θx* + (1 − *θ*)*y*), are smaller or equal to a linear line between *f* (*x*) and *f* (*y*), i.e. *θf* (*x*) + (1 − *θ*)*f* (*y*) [25].

A quasi-convex function is a little different as the function value of any point *θx* + (1− *θ*)*y* between any other two *x, y* ∈*C* may not be larger than the maximum of the function values *f* (*x*) and *f* (*y*). Mathematically speaking,

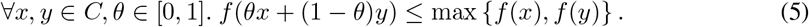

Therefore, every convex function is also quasi-convex, but not vice versa [25].

#### 1.2.3 Disciplined Convex Programmingming (DCP)

Disciplined Convex Programmingming is a methodology for constructing convex optimization models. The term “disciplined” emphasizes that DCP follows specific rules when constructing programs.

DCPs are motivated by the fact that it is theoretically and practically intractable to prove whether a particular non-linear program is convex. The idea of DCP is to offer an *atom library*, which is “an extensible collection of functions and sets, or *atoms*, whose properties of curvature/shape (convex/concave/affine), monotonicity, and range are explicitly declared” [26, p.171] and a *convexity ruleset* that defines “how atoms, variables, parameters, and numeric values can be combined to produce convex results” [26, p.171]. If an expression is constructed using only methods from DCP, it can be inferred which properties that expression has. For example, let *X* be a variable, then *Y* = *X* ·*X* is not valid under DCP because one cannot guarantee that the product of two variables is convex or concave for any two variables. But *Y* = POW(*X*, 2) is valid under DCP because, for the power function, one can guarantee the properties of the result.

Following Agrawal and Boyd [27], DQCP is an analog of DCP for quasi-convex functions.

### 1.3 Binary Linear Programming

Given a set of data points or entities, we aim to assign them to splits meeting some constraints, and we will use Binary Linear Programming formulated using DQCP rules to this end. The underlying idea is to assign a binary variable to any pair of entities and split that defines whether that entity is assigned to that split. We will explore this more in section 2.2. *Binary Linear Programs* are conceptually very similar to well-known *Integer Linear Programs*. Whereas the latter are linear optimization tasks with the restriction that the optimized variables have to be integers, in Binary Linear Programming, the variables are constrained to 0 and 1.

### 1.4 Previous Work

In machine learning, the problem of Information Leakage has been acknowledged. Some tools, such as TDCommons[15], RDKit[28], or DeepChem[16], offer complex data splits as a part of a broader framework. As TDCommons (as in PyTDC v0.4.1 in PyPI) and the RDKit offer a subset of splits that DeepChem v2.7 provides, we will only discuss the splits implemented in DeepChem. In DeepChem, splitting methods can be divided into random splits, user-defined splits, or cold-drug splits that use clustering. The user assigns samples to splits for the user-defined splits, and DeepChem follows those instructions. Thus, user-defined splits will not be discussed further. Recently, two new methods have been published that are similar to the idea of DataSAIL but not as comprehensive, namely the LoHi-Splitter[17] and GraphPart[18]. In the following, we will briefly describe the methods we compare to in Section 4. All of them can split data only in one dimension.

#### DeepChem - Butina

This mode is based on the clustering algorithm by Butina[29]. After computing clusters with the Butina algorithm, they are assigned into splits starting from the largest clusters. This does not take into account inter-cluster similarities.

#### DeepChem - Fingerprint

In this mode, one can only split data into two sets simultaneously. DeepChem computes all pairwise Tanimoto scores between the fingerprints of the input molecules and assigns a random molecule to the first split. The remaining molecules are iteratively assigned by calculating the most dissimilar molecule to one of the two splits and assigning it to the other.

#### DeepChem - MaxMin

The MaxMin splitting is an application of the Maximum Minimum Diversity Problem described in Porumbel et al. [30]. This algorithm maximizes the minimum diversity between data points in different partitions of a dataset, e.g., molecules in different data splits.

#### DeepChem - Scaffold

In this procedure, the molecules are first clustered based on their Bemis-Murcko Scaffold[31], then the clusters are assigned to splits, starting with the largest, as in the Butina splitting.

#### DeepChem - Weight

Splitting by weight in DeepChem works similarly to Scaffold splitting and Butina splitting. The molecules are sorted by molecular weight from heavy to light and assigned to the splits in that order. Therefore, all heavy molecules are in the first split, and all light molecules are in the last split.

#### LoHi Splitter

LoHi Splitter focuses on two machine learning settings on small molecules: one for application in Lead optimization (Lo) and one for Hit identification (Hi) tasks. Here, we only use the Hi splitting algorithm that defines ILPs to compute data splits. The basis is a graph with nodes for all molecules (or clusters of them) in the dataset. Edges between them represent a pairwise similarity above a user-defined threshold. Based on this graph, the algorithm constructs an ILP to solve the balances vertex minimum *k*-cut problem. The idea is to remove as few edges as possible to assign the resulting connected components into the splits matching the size constraints. A downside of this is that the algorithm does not always find a solution, and the user needs biological knowledge to define meaningful thresholds. It works only for small chemical molecules, even though it can easily be transferred to other data types.

#### GraphPart

In contrast to the LoHi Splitter, GraphPart is implemented to split DNA/RNA and protein sequences. It uses an iterative algorithm optimizing an initial clustering of the data to optimize an initial data clustering using MMseqs2 or the Needleman-Wunsch algorithm.

## 2 Theory

### 2.1 Quantification of Information Leakage

First, we discuss the mathematical foundation for DataSAIL. This includes quantifying Information Leakage and formulating the problem of splitting data as a graph problem and then translating this into Binary Linear Programs. This will help us understand the complexity of the topic and how DataSAIL optimizes the data split for a given dataset.

Kaufman et al. define Information Leakage qualitatively: either a setting leaks information or not [9]. Elangovan et al. define a quantitative measure for Information Leakage for NLP tasks [32]. There, they sum the maximal similarity per sample in the evaluation data to any sample in the training data. Divided by the number of samples, this leads to the average leak per test sample.

We find this is incomplete, as only the biggest leak per test sample is considered. Therefore, we redefine the amount of leaked information between two datasets *P* and *Q* as

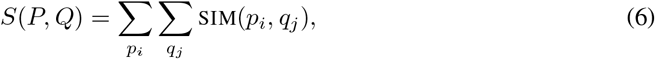

where SIM (·,·) is a function computing the pairwise similarity of the two inputs. With this, we can define the total pairwise similarity *S* of a dataset *D* with *n* data points *d* as

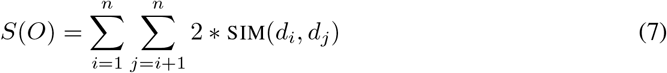

With this and partitioning *O* into *P* and *Q*, we can define *S*(*O*) (Figure 4A) as

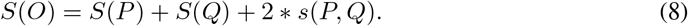

**Figure 4:**
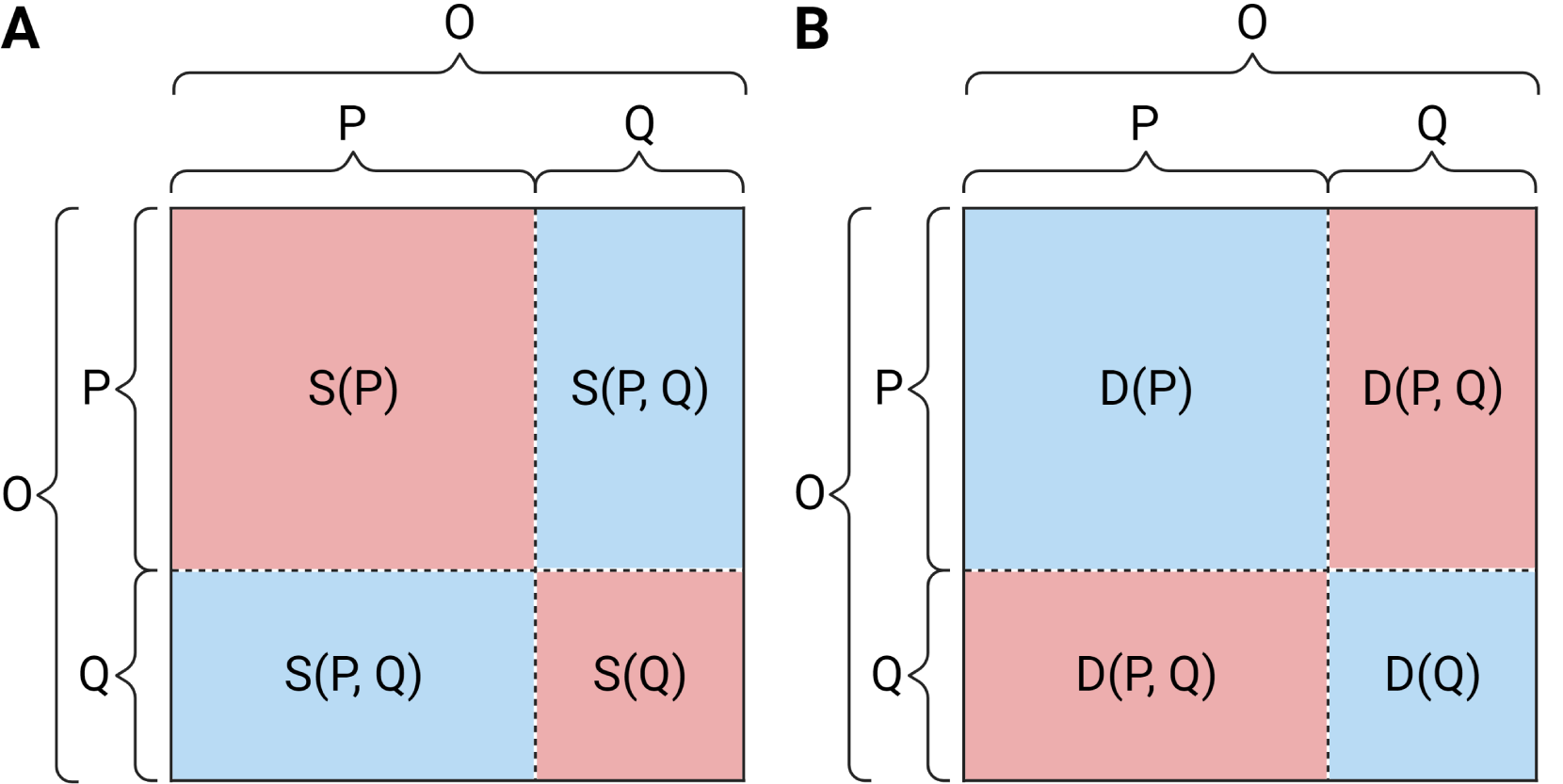
Visualization of Information Leakage based on a similarity measure (A) and a distance measure (B). The Information Leakage is lower in a data split with high intra-split similarity and a low inter-split similarity. From this, it follows that the inter-split distance has to be high and the intra-split distance low.

When no similarity metric is used, we define Information Leakage based on a distance metric. This can be done in the same way as in Equation 8 (Figure 4B):

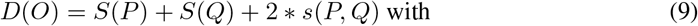

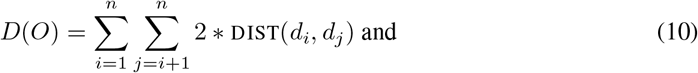

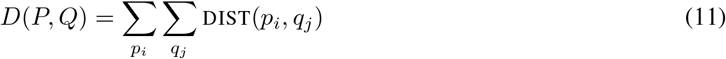

with DIST being the distance metric. Our reason for not converting distances into similarities is that there are multiple ways for conversion. Therefore, this would introduce an additional bias we want to prevent. From Figure 4 and Equations 8 and 9, minimizing the Information Leakage is equivalent to minimizing *S*(*P, Q*) or *D*(*P*) + *D*(*Q*), while maintaining a pre-defined ratio between *P* and *Q*.

### 2.2 Formulation as a Binary Linear Program

We now formulate the general problem of splitting an *R*-dimensional dataset into *k* splits mathematically to prove it is NP-hard and then as a Binary Linear Program. The problem consists of

- *R* sets of entities *V*_*r*_ with entity types *r* ∈ [*R*]
- Cardinalities *κ* : *V* → ℕ _≥1_ for all *u* ∈ *V*
- We write 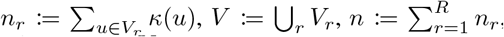, and let *t*(*u*) := *r* denote the type of an element *u* ∈ *V*_*r*_.
- Similarity scores *w*_*r,r*′_ : *V*_*r*_ × *V*_*r*′_ →ℝ_≥0_ for all unordered pairs of entity types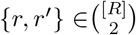
- Constants 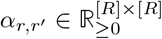 for all unordered pairs of entity types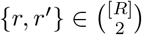
- We define *w* : *V* × *V* →ℝ_≥0_ as *w*(*u, v*) := *α*_*t*(*u*),*t*(*v*)_ · *w*_*t*(*u*),*t*(*v*)_(*u, v*).
- A desired number of splits *k* along with desired relative split sizes *s*_*i*_ ∈ (0, 1) with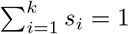
- An acceptable relative error *ϵ* ∈ [0, 1) for the relative split sizes.

We want to find a surjective mapping *π* : *V*→ [*k*] that splits *V* into *k* splits *π*_*i*_ := *π*^−1^[{*i*}] such that Information Leakage is minimized and the desired split sizes are preserved. For this, we define two objectives ([·] : {⊥, ⊺} → {0, 1} is the Iverson bracket):

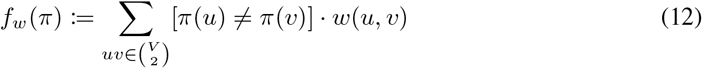

Moreover, we define constraints for every pair of dimensions and split:

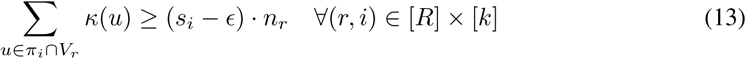

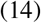

The DataSAIL problem can then be formulated as follows:

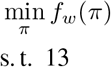

Based on this, we can formulate (*k, R*)-DataSAIL as a Binary Linear Program (BLP) as follows:

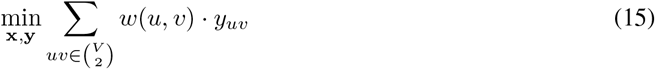

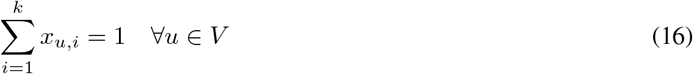

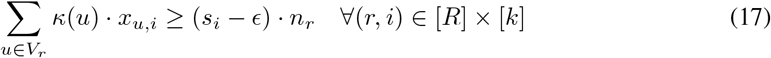

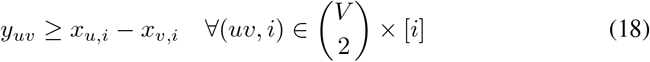

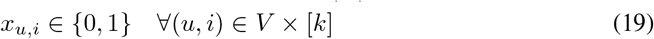

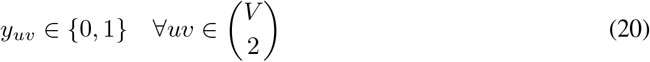

Constraints (16) and (17) ensure that **x** encodes a partition *π*_x_ satisfying constraint (13). Constraint (18) ensures that

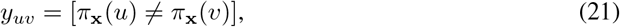

which implies that the objective minimized in (15) equals *f*_*w*_(*π*_x_). This, in turn, implies that the ILP given in the equations (15) to (20) is equivalent to (*k, R*)-DataSAIL. To see why (18) implies (21), note that the right-hand side of (18) is 0 for all *i* if *π*_x_(*u*) ≠ *π*_x_(*v*). Otherwise, the right-hand side of (18) is 1 for the cut *i* that contains *u* but not *v*. Since we minimize over *y* with non-negative coefficients in the objective, these considerations imply (21).

### 2.3 NP-hardness

We now show that the (*k, R*)-DataSAIL problem is *NP* -hard for all fixed constants *k* ∈ ℕ_≥2_ (number of splits) and *R* ∈ ℕ_≥1_ (number of entity types). We first show a polynomial time reduction from (*k*, 1)-DataSAIL to (*k, R*)-DataSAIL for arbitrary fixed *R* ≥ 2. Let 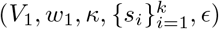 be an instance of (*k*, 1)-DataSAIL (we can ignore the *α* parameters for (*k*, 1)-DataSAIL). We now construct and instance of (*k, R*)-DataSAIL by adding *R* − 1 copies 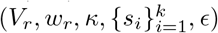 of 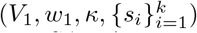. Let *u*_*r*_ denote the copy of node *u*_1_ ∈ *V*_1_ in copy *r* of the original (*k*, 1)-DataSAIL instance. We define *κ*(*u*_*r*_) := *κ*(*u*_1_) for all node copies, *w*_*r,r′*_ (*u*_*r*_, *v*_*r′*_) := *w*_1,1_(*u*_1_, *v*_1_) if *u ≠ v* or *r* = *r*^′^, and *w*_*r,r′*_ (*u*_*r*_, *u*_*r′*_) := *M* if *r*≠ *r*^′^, where *M* is some large enough constant 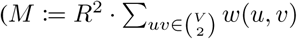 suffices). That is, edges between different copies of the same node get a very high weight *M*, and all other edges inherit their weights from the (*k*, 1)-DataSAIL instance.

Given an optimal solution *π*_1_ for our (*k*, 1)-DataSAIL instance, we can always define an induced solution *π*_*R*_ for the (*k*, 1)-DataSAIL as *π*_*R*_(*u*_*r*_) := *π*_1_(*u*_1_) (constraint (13) continues to hold by definition of *κ*). For each edge *u*_1_*v*_1_ contained in the cut induced by *π*_1_, there are 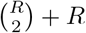 copies contained in the cut induced by *π*_*R*_, all of which have weight *w*(*u*_1_, *v*_2_): *R* copies of the form *u*_*r*_*v*_*r*_ and 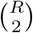copies of the form *u*_*r*_, *v*_*r′*_ with *r* ≠ *r*^′^. Moreover, the cut contains no other edges since, by definition of *π*_*R*_, all copies of the same node do not end up in the same split. Hence, we have

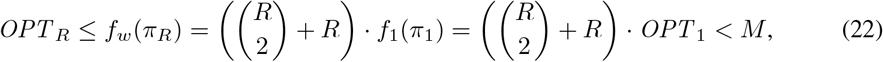

where *OPT* _1_ and *OPT* _*R*_ denote the optima of the (*k*, 1)-DataSAIL and the (*k, R*)-DataSAIL instances, respectively.

Conversely, let 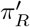 be an optimal solution for the (*k, R*)-DataSAIL instance. Then 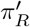 puts all copies of the same nodes into the same splits, since otherwise, we would have 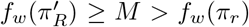, contradicting the optimality of 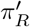. For all *u*_1_ ∈ *V*_1_, we now define 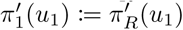. By counting edge copies as above, we obtain:

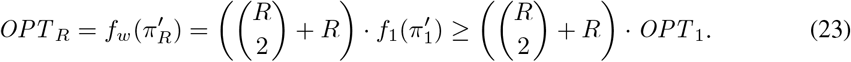

By combining the chains of inequalities (22) and (23), we obtain that 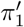 is optimal for our (*k*, 1)-DataSAIL instance, which finishes the reduction from (*k*, 1)-DataSAIL to (*k, R*)-DataSAIL.

It remains to be shown that (*k*, 1)-DataSAIL is *NP* -hard for all constants *k* ∈ ℕ _≥2_. This can be done via a straightforward reduction from the minimum *k*-section problem. Given a graph on *G* = (*V, E*) and a const *k* ∈ ℕ _≥2_, the minimum *k*-section problem asks to find a partition *π* into *k* splits minimizing Σ_*uv*∈*E*_[*π*(*u*) ≠ *π*(*v*)]such that

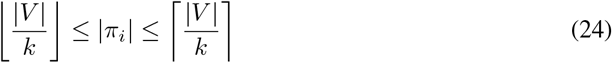

for all *i* ∈ [*k*] (all splits have the same size). This problem is *NP* -hard, even when restricting to balanced instances with |*V* | = *k* · *C* for some *C* ∈ ℕ_≥1_ [33], where the constraint (24) simplifies to

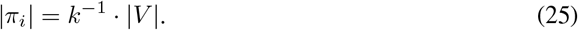

Given a balanced instance (*V, E, k*) of the minimum *k*-section problem, we now define an instance 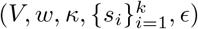 of (*k*, 1)-DataSAIL by setting *κ*(*u*) := 1, *w*(*u, v*) := [*uv* ∈ *E*], *s*_*i*_ := *k*^−1^, and *ϵ* = 0. Clearly, any solution *π* to (*V, E, k*) also solves the (*k*, 1)-DataSAIL instances and vice versa. Moreover, we have *f*_*w*_(*π*) = Σ_*uv*∈*E*_ [*π*(*u*) ≠*π*(*v*)] by construction of *w*. Consequently, solving (*V, E, k*) is equivalent to solving 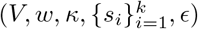, which completes the proof.

## 3 Methodology

After discussing the theoretical foundations of DataSAIL, we now focus on the implementation, the workflow, and the communication around the solvers. As mentioned in Section 1, DataSAIL primarily operates on biological macromolecules and splits them either based on identities or based on clusters (Figure 5).

**Figure 5:**
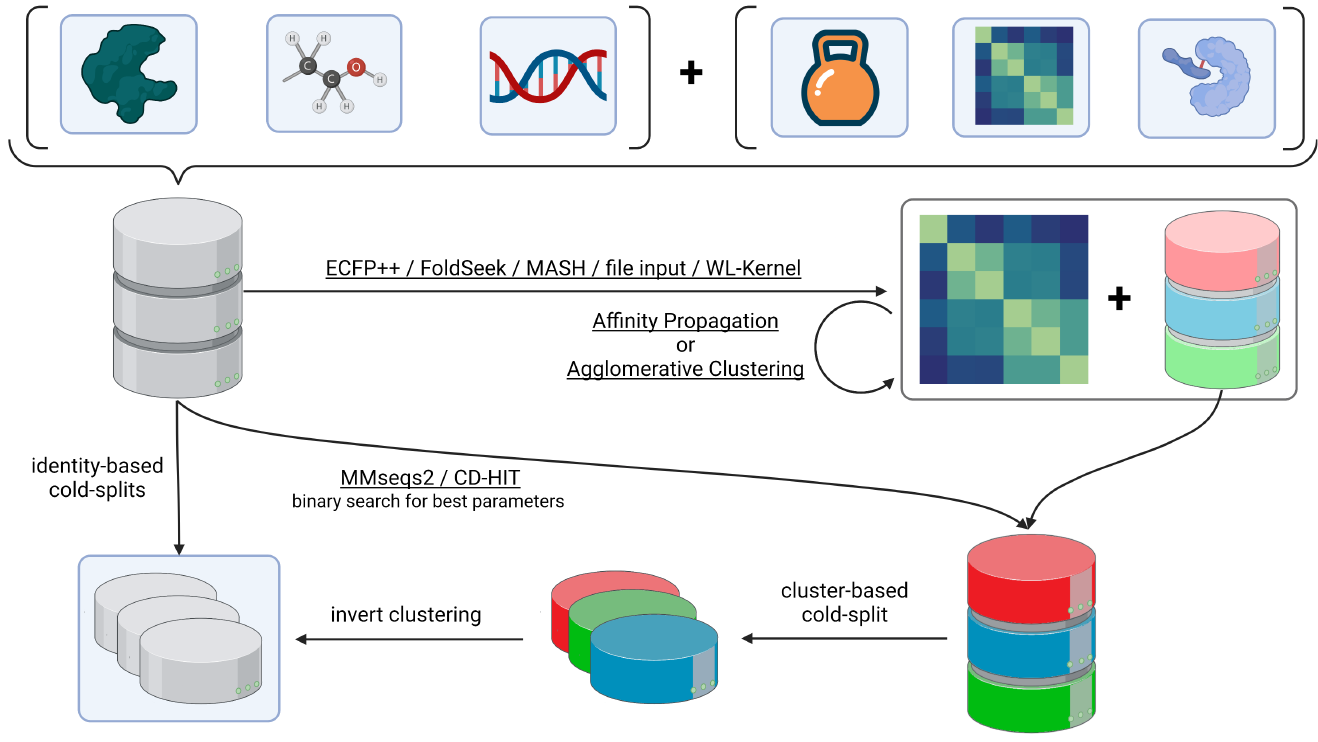
Graphical illustration of the workflow of DataSAIL. Input can be any data. We have clustering methods implemented within the pipeline for typical types of biomolecular data. Alternatively, user input can be supplemented with custom weighting, custom similarity or distance measures, and interactions between the input entities. Further input parameters may be specified. The output will be an assignment of input entities to splits. This figure was created with BioRender.com.

Identity-based splits are straightforward as the input is directly passed over to the solver as defined in Section 2.2. After solving, assignments of the entities to the different splits are extracted based on the values of *x*. For cluster-based splits, there are two options based on the input type. If the input is protein sequences, DataSAIL uses CD-HIT[34, 35] or MMseqs2[36] to produce a clustering of them. Both programs are not used to return a similarity matrix between the sequences but rather assign each sequence to a cluster; therefore, we computed a smaller instance of an identity-based split that is solved. Otherwise, DataSAIL computes or uses a user-defined pairwise similarity or distance matrix. To reduce the sizes of *x*, and ultimately the runtime and memory consumption of the solver, DataSAIL clusters the input data using spectral clustering[37] (for similarities) or agglomerative clustering[38] (for distances), as implemented in scikit-learn[39]. DataSAIL clusters the data down to 50 clusters. This number is picked as it leads to smaller problem instances to be solved but is still big enough to grasp some structure in the dataset.

It is important to note that the results of DataSAIL are non-deterministic by default and depend on the input order of the data. This is because the clustering algorithms may return results depending on the input data’s order and the solvers’ non-deterministic nature: multiple equally optimal solutions may exist, or the solver does not find the optimal solution within the time limits. To account for this, DataSAIL can run the solvers multiple times and get different splits for model evaluation.

DataSAIL is implemented as a command-line tool, and a Python package is installable from Conda[40]. Detailed information about the input parameters and examples of how to apply DataSAIL to exemplary datasets can be found on GitHub (https://github.com/kalininalab/DataSAIL) and ReadTheDocs (https://datasail.readthedocs.io).

## 4 Experiments

In the previous sections, we discussed DataSAIL’s approach to Information Leakage from a theoretical point of view. Here, we experimentally validate the idea and demonstrate the power of DataSAIL. To this end, we compute splits for various biological datasets using DataSAIL. Then, to assess to which extent information is leaked across splits, we train a machine-learning model from a classical collection and compare different splitting techniques regarding the model’s performance. It is expected that if Information Leakage occurs during training, the performance will be overestimated for identity-based splits and drop for cluster-based splits. DataSAIL can split one-dimensional and two-dimensional data, so we consider datasets and models for both settings.

To assess DataSAIL for splitting one-dimensional molecular data, we use the MoleculeNet datasets [21], a widely used collection of datasets representing properties of small molecules from various studies in quantum mechanics, physical chemistry, biophysics, and physiology (Table 1).

**Table 1:**
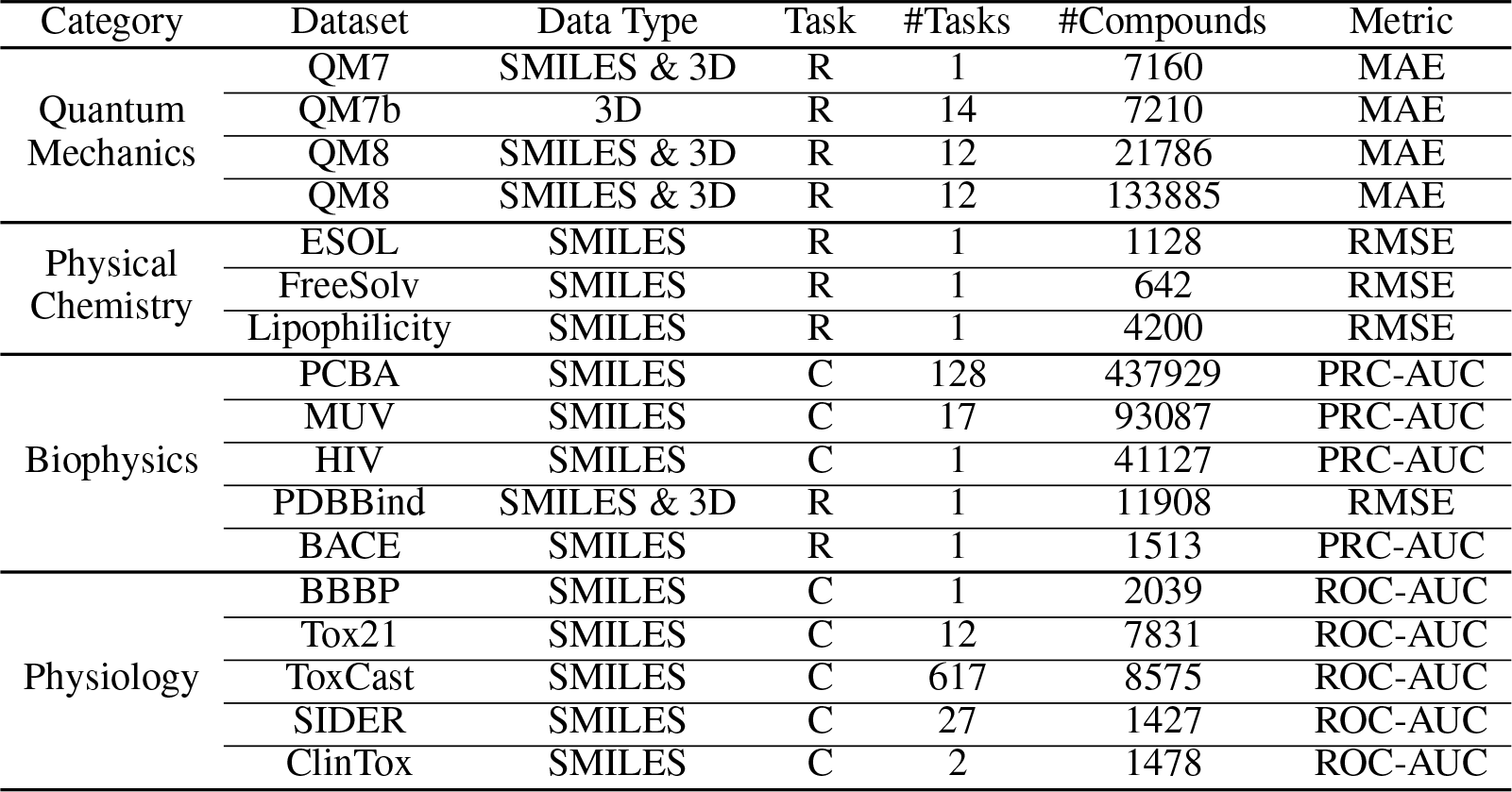
Overview of the datasets in MoleculeNet.

### 4.1 One-dimensional datasets

In the following, we discuss the performance of splitting algorithms on different MoleculeNet datasets in detail. Figure 6 shows all tSNE embeddings of the molecules labeled with their respective splits per method. One can see that the compactness of orange dots (test class molecules) differs heavily between datasets and splits. Sometimes, e.g., in the LoHi split of QM8 or fingerprint split of QM9, there is a clear structure where the test set has been clearly separated from the training set. But the same techniques compute less structured splits, for example, for FreeSolv, where other techniques compute better splits based on the 2D visualization. This indicates that there is not one best splitting method for all datasets, but the choice of the method should depend on the dataset. This is backed by Figure 7, where splits that lead to complex splits for QM9 pose easier challenges on FreeSolv.

**Figure 6:**
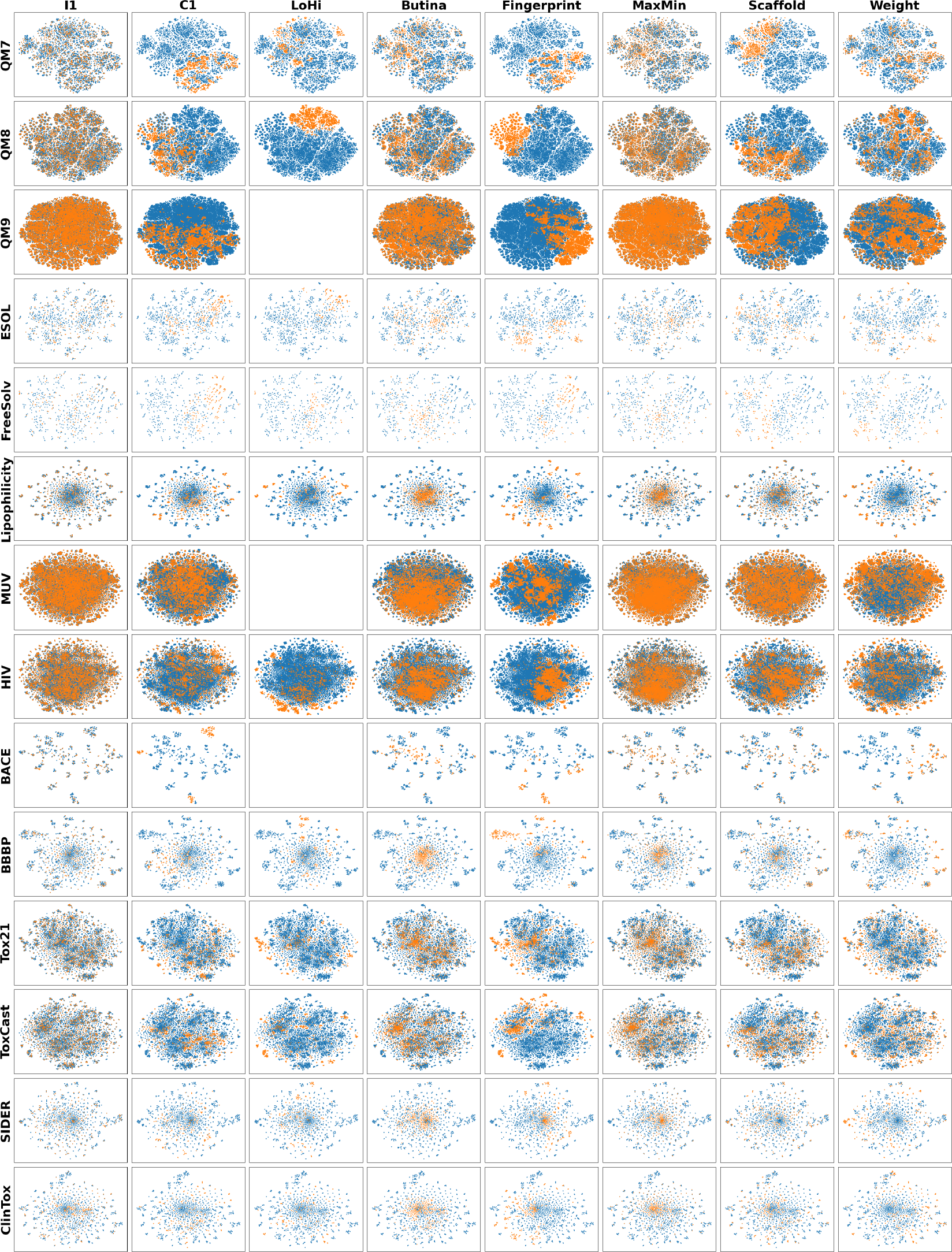
t-SNE embeddings of the molecules in the individual datasets of MoleculeNet. The embeddings were computed based on the 1024-bit ECFP4 fingerprints of the molecules using radius 2. The two left columns are DataSAIL’s splits (I1 is identity-based and C1 is cluster-based), the third is the LoHi splitter, and the five columns on the right are DeepChem’s algorithms.

**Figure 7:**
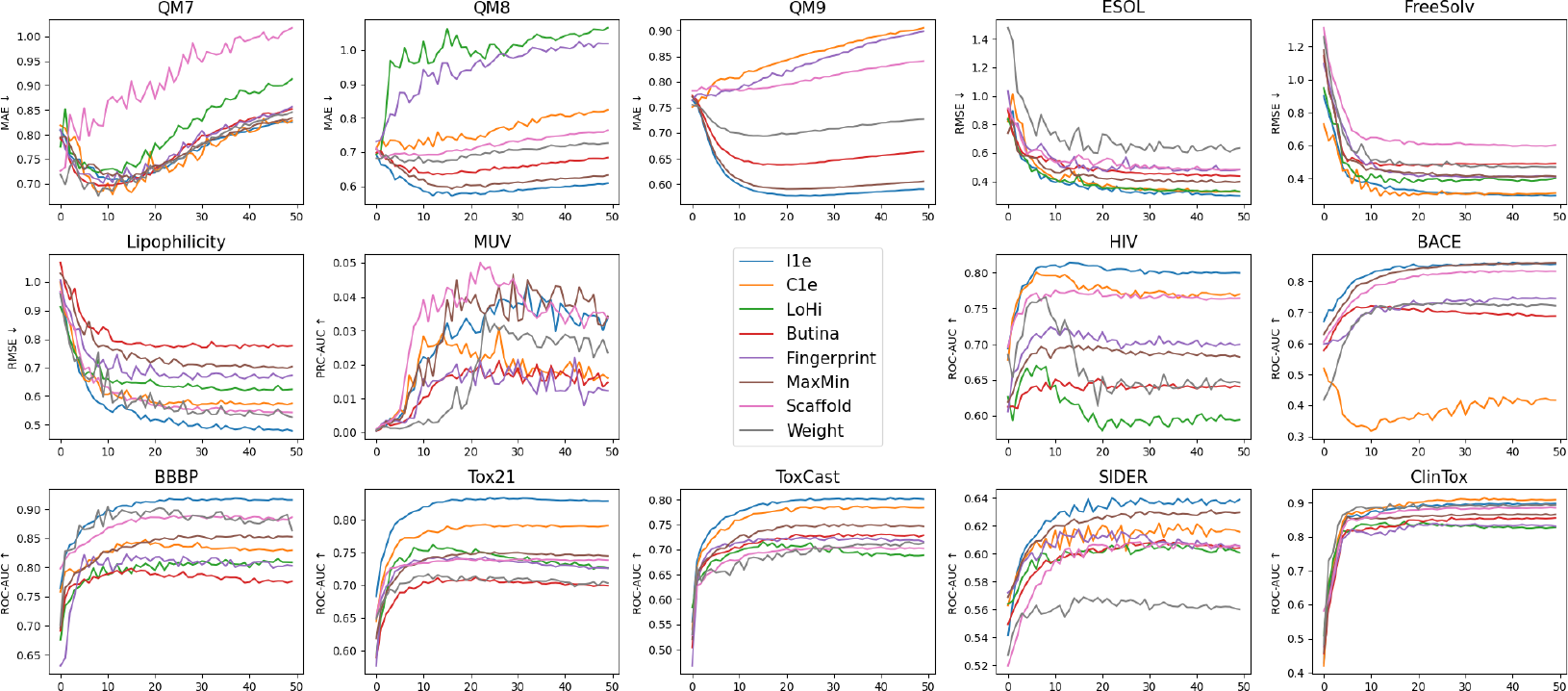
Performance comparison of different splitting algorithms.

Another observation is that not all plots are full. As discussed in Section 1.4, the LoHi splitter is slow and consumes much memory. When splitting MUV and QM9, the process was terminated after eight hours due to excessive memory usage. Furthermore, we do not show results for the PCBA dataset in MoleculeNet for the same reason. PCBA is too big to be split with any tool on our servers; it consists of ∼ 440k molecules.

#### 4.1.1 Random split vs cluster-based split on QM9 and Tox21 datasets

Here, we will look at the QM9 and Tox21 datasets in detail and show the impact of DataSAIL’s cluster-based split over a random split (here called *identity-based split*). In QM9, the task is to predict the quantum mechanical properties of molecules with up to nine heavy atoms. In Tox21, the task is to predict if a molecule is active or inactive in twelve different toxicological experiments.

When cluster-based DataSAIL splitting is used, the data points from the test set form distinct groups in the space of reduced dimensions, consistent with the procedure of cluster-based splitting (Figure 8, left and middle panels). Accordingly, the D-MPNN performance drops for this training regime, as the model is forced to generalize into these unseen areas of the data space instead of memorizing the neighboring points, which is the definition of reducing the Information Leakage (Figure 8, right panels). From this insight and Figure 7, we can see that our cluster-based splits pose challenging splits on the model testing for its ability to generalize to unseen molecules.

**Figure 8:**
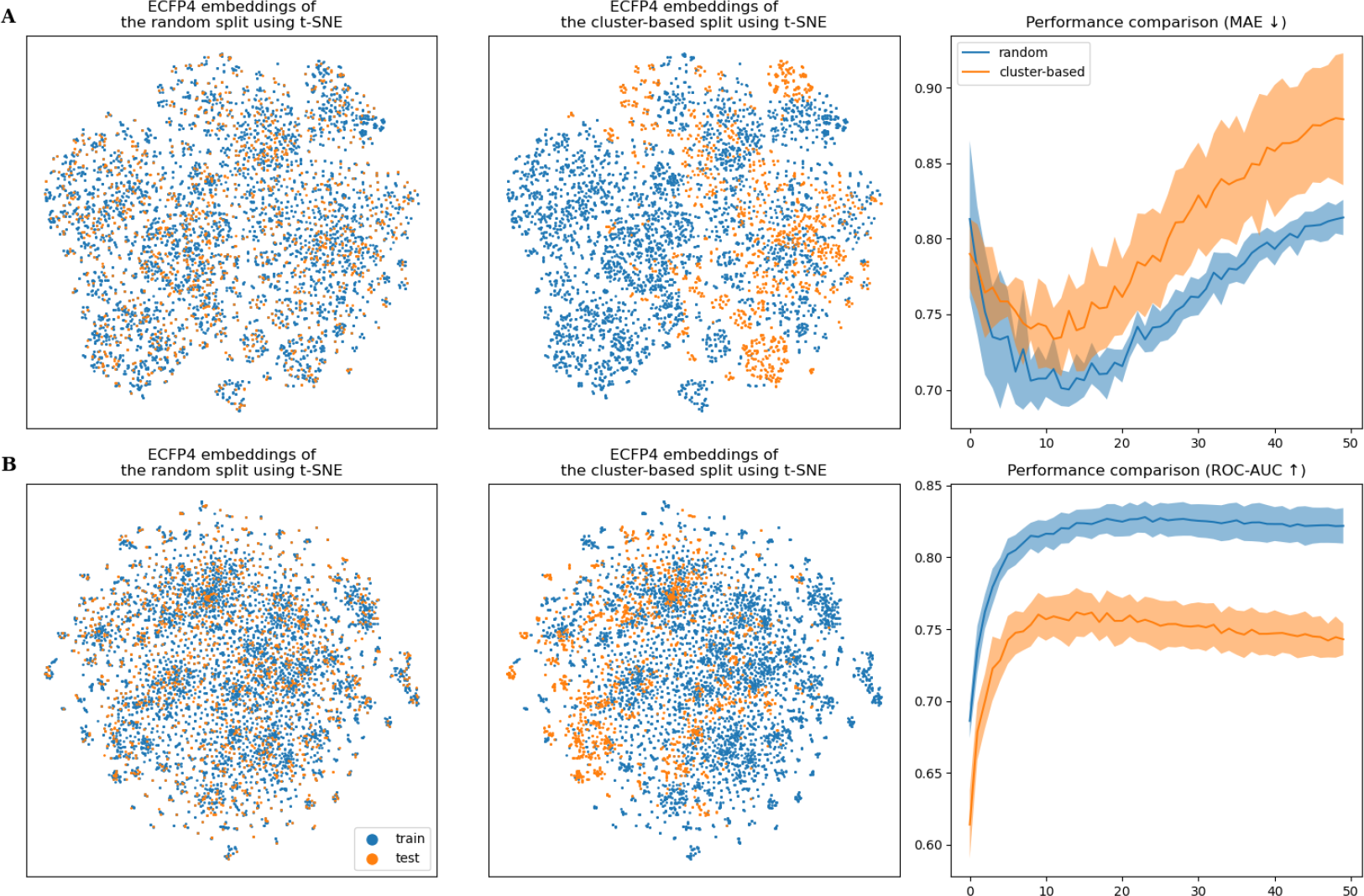
Visual analysis of the QM9 (A) and Tox21 (B) datasets. Left and middle: t-SNE embeddings of the ECFP4 fingerprints of the molecules in a random split (left) and a cluster-based split (middle). Right: Performance estimate in the training of D-MPNN, solid lines show averages over the performances on five different splits computed with the same technique, and the shaded area shows their standard deviation.

#### 4.1.2 Runtime

Because MoleculeNet offers a variety of datasets with different sizes, structures, and similarities, we use it for benchmarking the runtime of the splitting tools(DataSAIL, LoHi splitter, and DeepChem, Figure 9).

**Figure 9:**
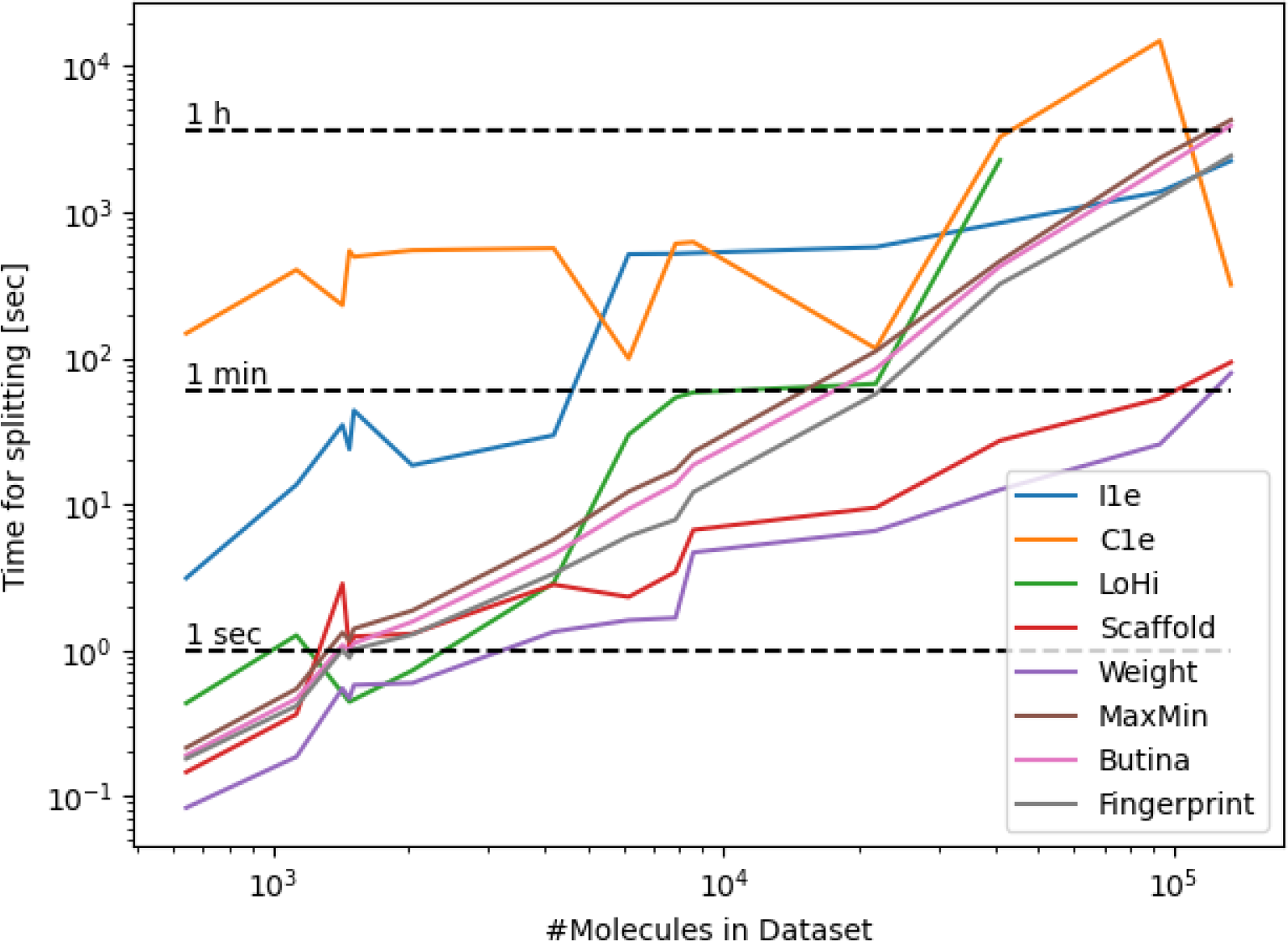
Runtime analysis on MoleculeNet. Both axes have been log-transformed to improve the interpretability of the results.

As expected, the bigger the datasets, the slower the algorithms compute their splits. However, it is important to note that the LoHi splitter does not produce results for datasets above a certain size (see also Section 4.1). Furthermore, cluster-based splitting by DataSAIL scales much better than other algorithms because the problem size roughly stays the same due to the clustering step.

### 4.2 Two-dimensional case: LP-PDBBind

To evaluate DataSAIL on two-dimensional data, we use LP-PDBBind[41]. This is an improved dataset based on the PDBBind dataset combined with results from BindingDB. It contains a set of experimentally resolved three-dimensional structures of protein-ligand complexes. We split it with DataSAIL employing seven different techniques: a random split, identity-based and cluster-based splits along each axes (proteins and ligands), and a cluster-based two-dimensional split.

The model we use for training on these splits is DeepDTA[20], which aims to predict whether a protein binds a given ligand or not. It achieved top performances on the state-of-the-art KIBA[42] and Davis[43] datasets and is easy to use[44]. DeepDTA uses sequence representation for proteins and SMILES for ligands.

In this experiment, identity-based splits (both for proteins and drugs) lead to a model performance that is very close to a random split. Again, in training with the cluster-based splits, the model performance drops dramatically, and the cluster-based split along the protein axis is as detrimental or even worse for the model performance as the cluster-based double split (Figure 10). This indicates that the Information Leakage along the protein axis is stronger than along the drug axis, which is also evident when analyzing the distribution of the data points in the data space (Figure 10).

**Figure 10:**
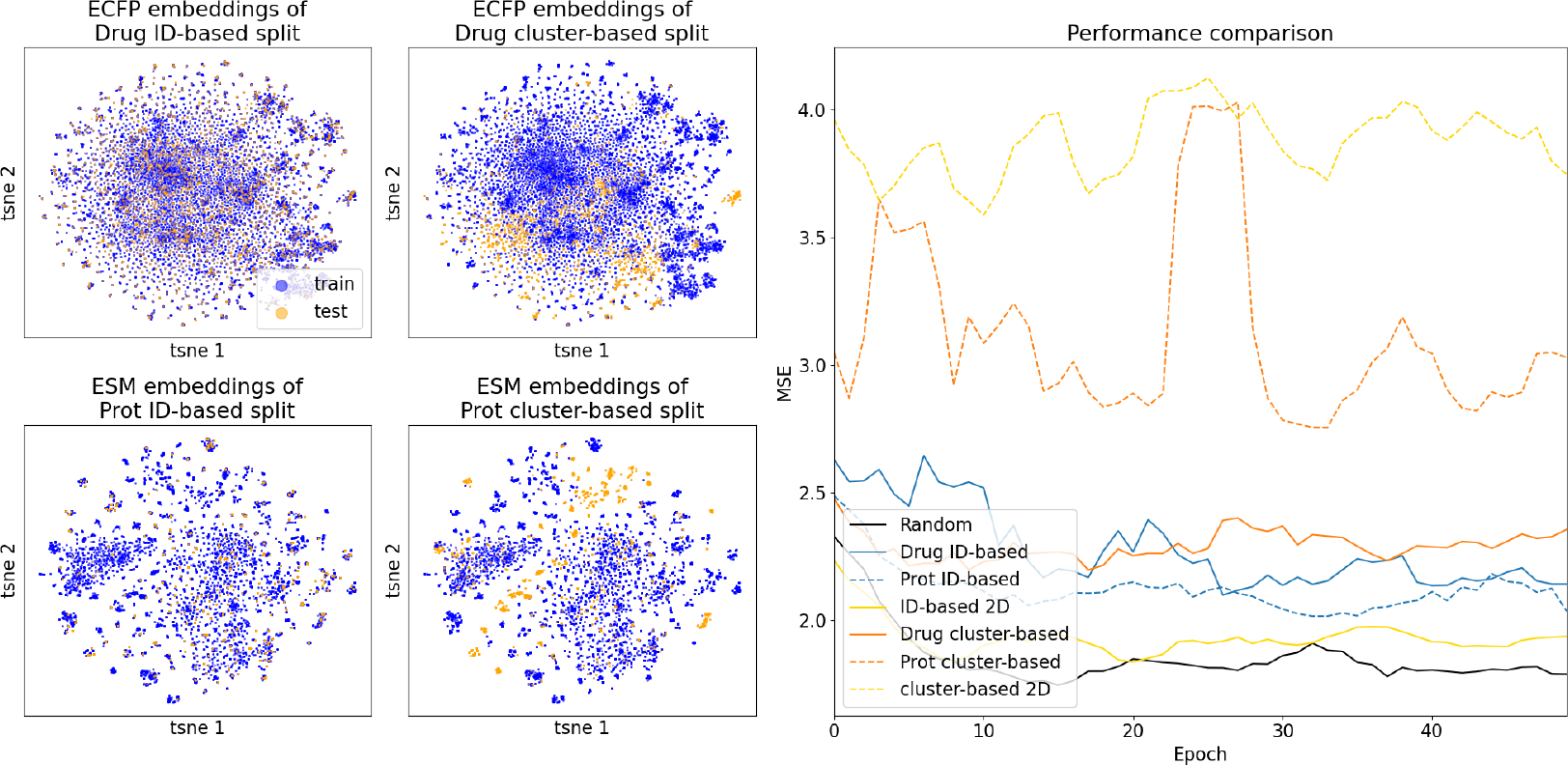
Visual analysis of LP-PDBBind. Left panel: t-SNE embeddings of ligands (upper plots) and proteins (lower plots). Right panel: Comparison of seven different splitting techniques on LP-PDBBind. The lines are averaged over five attempts to split the dataset with the same technique and smoothed using the floating window method with a window size of 5.

#### 4.2.1 One-dimensional splits of LP-PDBBind under the microscope

To our knowledge, DataSAIL is the only tool splitting two-dimensional data along both dimensions simultaneously. But there are settings where splitting along one axis is best because,e.g., the model will only see new drugs but no new proteins at the inference time. This is the case in drug development for a known, fixed target protein, e.g., against the spike protein of the COVID-19 virus [45]. Therefore, we will look at how different splitting algorithms perform on LP-PDBBind when splitting only drugs or targets.

We compare DataSAIL with DeepChem and LoHi splitters for drugs (Figure 11). The LoHi splitter performs best in creating a tough split for the model, supported by the visual analysis of the separation between training and test sets in the data space.

**Figure 11:**
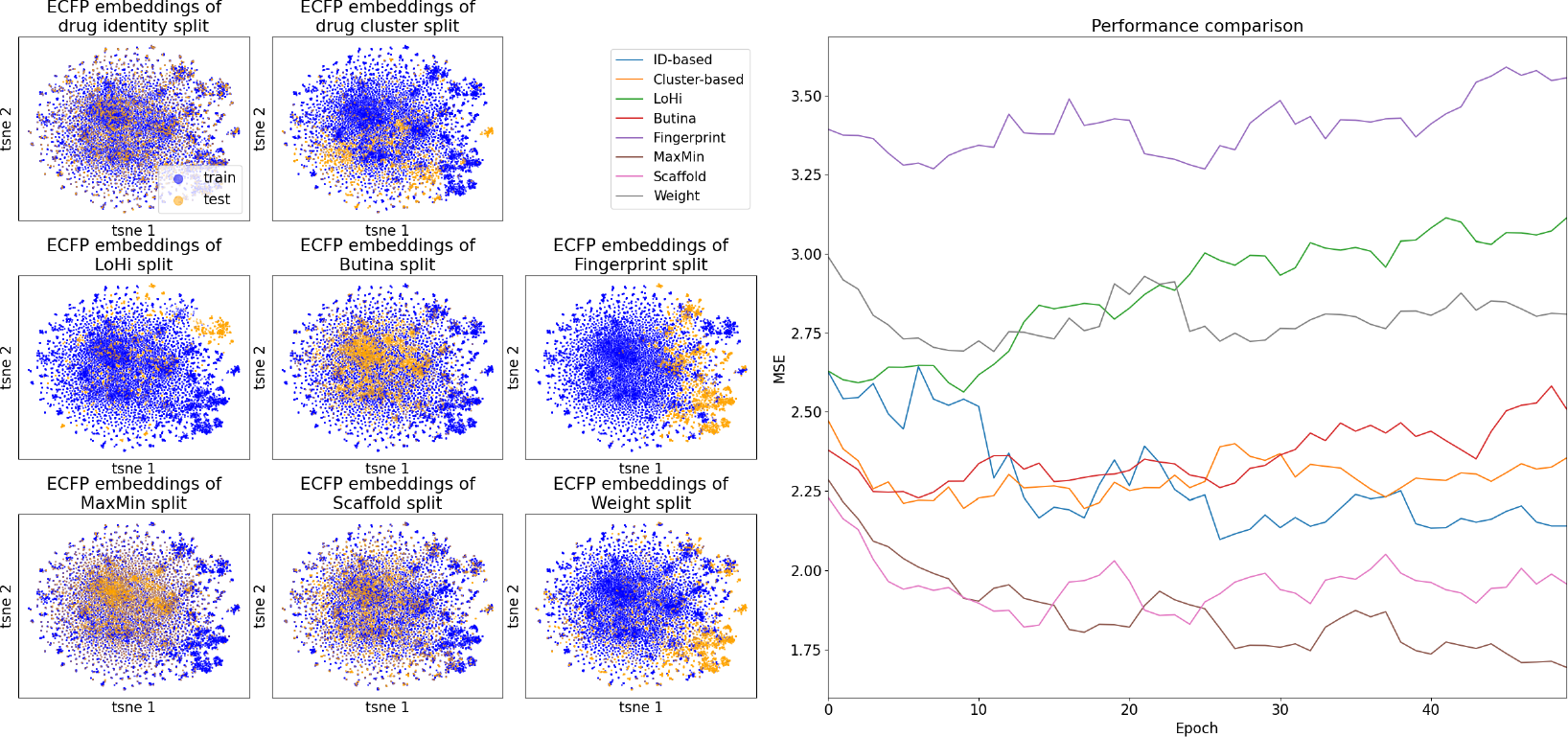
Comparison of splitting methods for molecule dataset of LP-PDBBind.The lines are averaged over five splits of the dataset with the same technique and smoothed using the floating window method with a window size of 5.

For target proteins, we can compare DataSAIL only with GraphPart. Both GraphPart and DataSAIL cluster-based split lead to a separation between training and test data (Figure 12, left panels), and they are equally difficult for the model to generalize (Figure 12, right panel).

**Figure 12:**
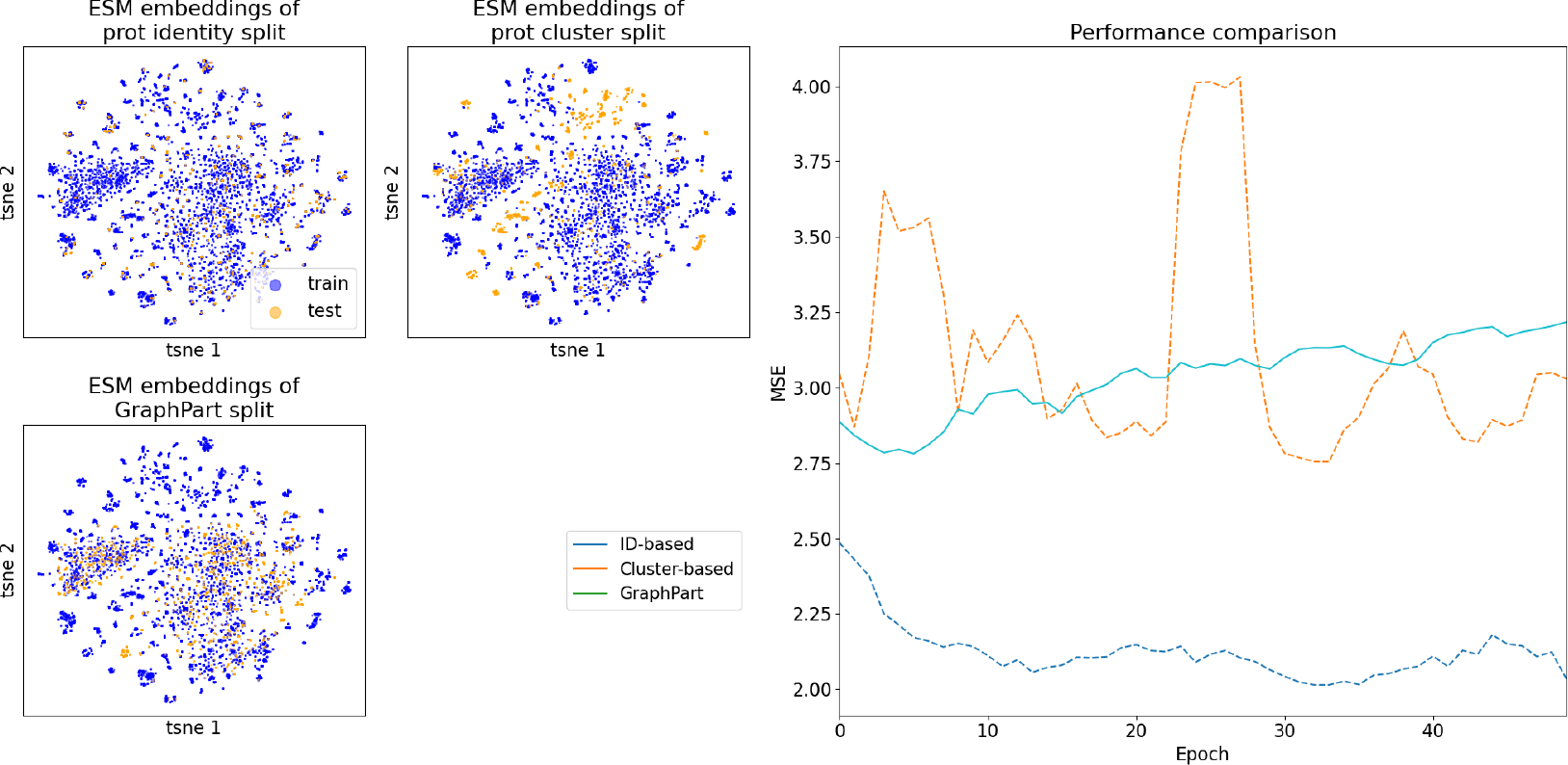
Comparison of splitting methods for protein dataset of LP-PDBBind. The lines are averaged over five splits of the dataset with the same technique and smoothed using the floating window method with a window size of 5.

## 5 Conclusions

This study discusses the importance of Information Leakage in training machine-learning models for biomedical research. Whereas Information Leakage is a general term with many facets [10], we focus on the case when a too high similarity between training and test sets causes the model to memorize the data instead of extracting predictive properties. Not accounting or improperly accounting for Information Leakage can lead to inflated reported performance of the models and incorrect conclusions, as demonstrated in the experiments.

We propose a computational strategy and a tool, DataSAIL, to minimize such Information Leakage. We formulate data splitting as a set of binary linear optimization problems, prove its NP-hardness, and offer an efficient computational solution for optimizing them. We offer one framework to split one-dimensional and two-dimensional data (when data can be represented as an array or a matrix). Whereas we focus on molecular data (small molecules and protein sequences), our framework can easily accommodate other data types, provided one can calculate the similarity or distance between the data points.

We apply DataSAIL in several scenarios relevant to predicting the properties of small molecules and drug-target interactions and demonstrate a distinct drop in performance when Information Leakage is removed. Further, the proposed framework can be readily applied in other biological problems where machine learning is routinely used, for example, the prediction of mutation effects or protein-protein interactions.

Compared to DeepChem, the LoHi splitter, and GraphPart, we can see that DataSAIL provides more challenging splits for the models in a reasonable time. It is the first tool reducing Information Leakage in two-dimensional datasets on both dimensions simultaneously. Unlike other tools, when applied to one-dimensional data, DataSAIL does not remove any data from splits. DataSAIL can compute multiple splits simultaneously without reloading the data. Furthermore, it always computes a solution for a valid input and does not require domain knowledge. Additionally, DataSAIL is faster on larger datasets because it scales better to large amounts of data. To sum up, DataSAIL is a Swiss army knife for data splitting, providing all kinds of data splits with one line of Python code.

